# Geobacter sulfurreducens’ unique metabolism results in cells with a high iron and lipid content

**DOI:** 10.1101/2022.07.29.502083

**Authors:** Ethan Howley, Dongwon Ki, Rosa Krajmalnik-Brown, César I. Torres

## Abstract

*Geobacter sulfurreducens* is a ubiquitous iron reducing bacterium in soils, and in engineered systems it can respire an electrode to produce measurable electric current. Its unique metabolism, heavily dependent on an extensive network of cytochromes, requires a unique cell composition. In this work we used metallomics, cell fraction and elemental analyses, and transcriptomics to study and analyze the cell composition of *G. sulfurreducens*. Elemental composition studies (C,H,O,N, ash content) showed a high C:O and H:O ratios of approximately 1.7:1 and 0.25:1, indicative of more reduced cell composition that is consistent with a high lipid content. Our study shows that *G. sulfurreducens* cells have a large amount of iron (2 ± 0.2 μg/gdw) and lipids (32 ± 0.5% dw/dw) and that this composition does not change whether the cells are grown with a soluble or an insoluble electron acceptor. The high iron concentration, higher than similar microorganisms, is attributed to the production of cytochromes that are abundant in transcriptomic analyses in both solid and soluble electron acceptor growth. The unique cell composition of *G. sulfurreducens* must be considered when growing this microorganism for lab studies and commercial applications.

**Importance:** *Geobacter sulfurreducens* is an electroactive microorganism. In nature, it grows on metallic minerals by transferring electrons to them, effectively ‘breathing’ metals. In a manmade system, it respires an electrode to produce an electric current. It has become a model organism for the study of electroactive organisms. There are potential biotechnological applications of an organism that can bridge the gap between biology and electrical signal, and as a ubiquitous iron reducer in soils around the world, *G. sulfurreducens* and its relatives impact the global iron cycle. We measured the concentrations of metals, macromolecules, and basic elements in *G. sulfurreducens* to define this organism’s composition. We also used gene expression data to discuss which proteins those metals could be associated with. We found that *G. sulfurreducens* has a large amount of lipid and iron compared to other bacteria — these observations are important for future microbiologists and biotechnologists working with the organism.

## Introduction

Anode-respiring electroactive bacteria, such as *Geobacter sulfurreducens*, have been studied for almost two decades for their capability to produce electrical current from metabolic respiration of organic compounds while in multi-layered biofilms (1–3). A unique feature of these biofilms is the extracellular matrix that allows the transport of electrons over tens of micrometers (4–6). As part of this extracellular matrix, several components have been proposed to be crucial in achieving extracellular electron transport (EET). The transport of electrons starts at the inner membrane, and travels across the periplasm and outer membrane before it reaches the extracellular environment. Cytochromes at these locations are known to play an important role in delivering electrons outside the cell (7–10). Microbial nanowires, now also identified as cytochrome polymers (4, 6), are the main path by which electrons are thought to be conducted in the extracellular environment, reaching a solid electron acceptor. Extracellular polymeric substances (EPS) have also been proposed to play a role in EET in *G. sulfurreducens* as well (3, 11). On the other hand, *Shewanella oneidensis* MR-1 has been shown to produce outer membrane and periplasmic extensions, which are lipid bilayers and contain extracellular cytochromes (12, 13). In both cases, it is clear that the EET mechanism creates an extra metabolic burden to electroactive organisms and that it can alter their cell composition and nutrient requirements when compared against microorganisms performing respiration of soluble electron acceptors. These nutrient requirements, however, have not been assessed in a systematic way.

Transcriptomic and proteomic studies in *G. sulfurreducens* have highlighted the importance of respiratory and EET proteins for their growth on anodes and metal oxides. Extracellular and outer membrane proteins, including pili, outer membrane channels, and c-type membrane cytochromes, are essential to the metabolism of *G. sulfurreducens* (9, 14–26). The high abundance of these proteins and other possible extracellular components may result in a unique cellular composition. For example, each cytochrome contains one or more iron-containing heme complexes which can influence the iron content of the cell. An analysis of cell composition can provide insights into the composition of the extracellular matrix and EET-related components.

Several studies have provided insights into the cellular composition of *G. sulfurreducens*.For example, the lipid fraction of *Geobacter sulfurreducens* was reported 15% wt/wt by Mahadevan et al., 2006, where an additional 4% lipopolysaccharides fraction was assumed (27). In comparison, the lipid in *Escherichia coli* has been reported to be 9.1% wt (28) while cyanobacterium *Synechocystis* sp. PCC6803, known to produce thylakoid membranes, has been reported to be as high as 14% wt(29) and the lipid-rich microalgae *Schizochytrium* sp. can contain up to 30% lipid (30). The high lipid content of *G. sulfurreducens* has potential implications for biotechnology applications.

To our knowledge, few studies have performed a metallomic analysis in *G. sulfurreducens*, and the existing literature has conflicting results. Previous research showed that *G. sulfurreducens*, when grown on fumarate as electron acceptor, has similar metal content to that of *E. coli*. On the other hand, the closely related organism *Geobacter metallireducens* grown on iron citrate showed an order of magnitude higher iron content; but the possible formation of inorganic precipitates was reported to be a possible hindrance to the measurement (31). Another study found that *G. sulfurreducens* had a per cell iron content an order of magnitude higher than *E. coli*, and that limiting growth medium iron content inhibited EET in *G. sulfurreducens* (15).

In this study, we hypothesized that *G. sulfurreducens* has a significantly different cellular composition compared to other cells performing soluble respiratory metabolisms. The differences that stem out of the EET requirements can help explain how EET develops. Understanding these characteristics can lead to a better growth and maintenance of this microorganisms in laboratory and applied systems. We performed a metallomic analysis on *G. sulfurreducens* grown on an anode versus fumarate as electron acceptors and compared it to *E. coli* K12. The use of an anode allows us to study EET and eliminates the interference of possible iron oxides reported by Budhraja et al. (31). We also performed an elemental analysis (C, H, O, N, ash content) and a fraction analysis (protein, carbohydrates, and proteins) to obtain a comprehensive cell composition and determine if the previously observed lipid fractions measured in fumarate samples are also observed during anodic respiration. The results are complemented with a transcriptomic analysis of *G. sulfurreducens* grown under similar conditions.

## Materials and Methods

### Bacterial strain and culture media

We subcultured *Geobacter sulfurreducens* PCA (ATCC 51573) and *Escherichia coli* K-12 from commercially available stocks. Medium compositions are listed in detail in Table S1 for four different cases: 1. *G. sulfurreducens* grown in microbial electrochemical cell (electrode), 2. *G. sulfurreducens* grown in sodium fumarate-containing serum bottle (fumarate), 3. *E. coli* grown in *Geobacter* medium, and 4. *E. coli* grown in M9 medium. In brief, *Geobacter* medium contained sodium acetate (50 mM), NaHCO_3_ (30 mM), NH_4_Cl (20 mM), NaH_2_PO_4_ (4 mM), KCl (1 mM), vitamin mix (10 mL), and trace minerals (10 mL). Trace minerals contained Nitrilotriacetic acid, trisodium salt (5.5 mM), MgSO_4_·7H_2_O (12 mM), MnSO_4·_H_2_O (2.9 mM), NaCl (17 mM), FeSO_4_·7H_2_O (0.36 mM), CaCl_2_·2H_2_O (0.68 mM), CoCl_2_·6H_2_O (0.42 mM), ZnCl_2_ (0.95 mM), CuSO_4_·5H_2_O (0.4 mM), AlK(SO_4_)_2_·12H_2_O (0.2 mM), H_3_BO_4_ (0.16 mM), Na_2_MoO_4_·H_2_O (0.01 mM), NiCl_2_·6H_2_O (0.01 mM), and Na_2_WO_4_·2H_2_O (8.5 μM). We provided higher concentration of acetate (50 mM) than the previous studies(32–34) (10 mM), as we used a higher electrode surface area requiring more reduced electron donor. For *G. sulfurreducens* grown in serum bottles, sodium fumarate (100 mM) was added in the medium. We bubbled the media with N_2_/CO_2_ (80:20 v/v) to remove oxygen before autoclaving. After autoclaving, FeCl_2_·4H_2_O (20 μM), Na_2_S·9H_2_O (54 μM), sodium bicarbonate, and vitamins were added in anaerobic glove box. *E. coli* medium (ATCC Medium 2511 - M9 Minimal Broth) contained glucose (44 mM), Na_2_HPO_4_ (180 mM), KH_2_PO_4_ (44 mM), NaCl (17 mM), NH_4_Cl (37 mM), MgSO_4_·7H_2_O (0.2 mM), CaCl2 (10 μM), and thiamin (24 μM). Media for *E. coli* (M9 and *Geobacter* medium) were autoclaved without any gas sparging. *Geobacter* medium for *E. coli* contained the same ingredients of *G. sulfurreducens* medium for microbial electrochemical cells except for the electron donor; the same concentration of glucose as M9 (44 mM) was used instead of acetate.

### Electrochemical setup and operation

Single-chamber microbial electrochemical cells were constructed in 500 mL bottles with rubber stoppers located on top having PTFE tubing for gas inflow and outflow, carbon anode as working electrode, nickel wire cathode as counter electrode, and reference electrode. We used two square graphite electrodes to grow biofilms of *G. sulfurreducens* with a surface area of ~20.9 cm^2^, and an Ag/AgCl reference electrode (BASi, West Lafayette, IN). We mixed the chambers with magnetic stirrer bars at 180 rpm and flushed humidified N_2_/CO_2_ gas (80:20 v/v) continuously. Before filling up the media in the anaerobic glove box, electrochemical cells were autoclaved for sterilization. We set −0.3 V vs Ag/AgCl (−0.03 V vs. SHE) as fixed anode potential using a VMP3 digital potentiostat (Bio-Logic USA, Knoxville, TN). This is at the lower end of the range of electrode potentials that *G. sulfurreducens* can respire. Fumarate-grown *G. sulfurreducens* reactors were set up in 250 mL serum bottles. Both electrochemical cells and serum bottles were in a temperature-controlled room at 30 °C. Also, the media filling along with inoculation was performed in the anaerobic glove box. We used 250 mL flasks for *E. coli* cultivation with M9 and *Geobacter* media. *Geobacter* medium (G medium) for *E. coli* was used to compare the cell composition with *G. sulfurreducens*. After filling the media and inoculation for *E. coli*, we placed the flasks in an incubator with shaking at 180 rpm and temperature at 37 °C.

### Sample preparation

For the determination of carbohydrate, protein, lipid, metal, and element per dried cell, we collected the *G. sulfurreducens* grown on anodes in the microbial electrochemical cells and in serum bottle reactors and the *E. coli* grown in flasks. *G. sulfurreducens* biofilms grown on the anodes were scraped off with a needle in an anaerobic glove box. Grown cells of *G. sulfurreducens* and *E. coli* from the serum bottles and flasks were separated by centrifugation (Eppendorf Centrifuge 5810 R, USA) at 4000 rpm in microcentrifuge tubes. Cells were washed once with a Ringer’s solution (25% strength) and centrifuged again (Table S1). The cells dried overnight at 105 °C in plastic tubes and the cool pellets were then broken up with a sterile stainless-steel spatula.

### Analytical methods

We measured proteins by bicinchoninic acid (BCA) protein assay (35). In brief, dried cell biomass (2-3 mg) treated with 0.1 N NaOH at 90 °C for 30 min, re-suspended and centrifuged the lysate, and used 0.1 mL of supernatant for the assay. Carbohydrates were measured by a colorimetric method (36). In brief, dried cell biomass (2-3 mg) was acidified in sulfuric acid with sonication for 2 hours, and dissolved samples (0.5 mL) added in the test tubes with distilled water (0.5 mL), phenol (50 μL), and sulfuric acid (5 mL) for overnight reaction. Concentrations of proteins and carbohydrates were determined using calibration curve with bovine serum albumin and glucose with the absorbance at wavelengths of 485 and 562 nm, respectively. Crude lipids were extracted from the dried cell biomass using the Folch method (37). The dried biomass (~15 mg) was sonicated for 1 hour and vortexed for 1 hour with Folch solution (chloroform-methanol, 2:1, v/v) at room temperature. Solvent extracts were obtained after removing the biomass by centrifugation at 4000 rpm. The crude lipid weight was determined by evaporating in the water bath (60 °C) and weighing the tube before and after the evaporation of lipid. For metal extraction, we added dried cell biomass (3-5 mg) to glass vials along with hydrochloric acid (12 M) and sonicated at 60 °C for 2 hours. The dissolved metals in acid were analyzed by inductively coupled plasma - optical emission spectrometer (ICP-OES, Thermo iCAP6300). Carbon, hydrogen, and nitrogen in the dried cell biomass (~2 mg) were measured using CHN Elemental Analyzer (PE2400). Instrumentation for ICP-OES and CHN Elemental Analyzer was done in Goldwater Environmental Laboratory at Arizona State University. For oxygen estimation, we measured ash content of the dried cell biomass (~10 mg) following the previous method (38), burning the biomass at 600 °C in an alumina crucible. Then we subtracted the fraction of C, H, N, and ash content from the dried cell biomass (100%) for O estimation. Based on the elemental analysis data (%C, %H, %O, and %N), we obtained the empirical biomass formula followed by Equation 1 below (39),

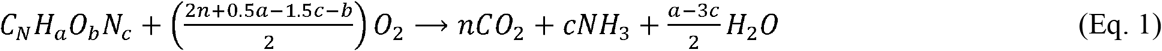

where, 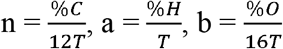, and 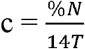,
and, 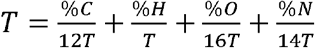.

### Transcriptomic analysis

We sequenced reverse transcribed RNA from *G. sulfurreducens* grown in a single chamber microbial electrochemical cell as an anode biofilm with an anode poised at −0.28 V vs. Ag/AgCl as the electron acceptor, and in anaerobic test tubes with fumarate as the electron acceptor. Both conditions were collected in biological triplicate. We extracted RNA with the Qiagen/MOBIO PowerMicrobiome RNA extraction kit and used the ThermoFisher MicrobExpress bacterial mRNA enrichment kit to reduce the fraction of rRNA following manufacturer’s recommendations. RNA library preparation and sequencing were performed by the genomics core facility at ASU. We mapped the reads to the RefSeq assembly for *G. sulfurreducens PCA* and used DESeq2 in R for differential expression analysis. The cutoff determination for differential expression was set as a log_2_ fold change of at least 1.5 and a multiple comparison adjusted *P*-value of less than 0.05. The sequencing data we used here is a subset of data analyzed in a previous publication, and raw sequence data is available from NCBI under accession number GSE200066 (40).

## Results and Discussion

### Elemental analysis

We analyzed the trace metals and macronutrients of dried *G. sulfurreducens* and *E. coli* grown in the lab with different environments (Table S3, Table 3, Figure 1). We found significant differences between the two species in the mass fraction of several trace metals (Figure 1, Table S3, Figure S1). Media compositions were different in each growth condition, matching each organism’s requirements (Table S2). There were few differences between anode-grown and fumarate grown *G. sulfurreducens*. Significant differences in Mn and Fe content were observed in *E. coli* when growing in M9 medium versus *Geobacter* medium. Given the low abundance of nutrients in M9’s minimal medium, significant increases in Cu, Fe, Mn, and Se were observed when *E. coli* was grown in *Geobacter* medium. We use this latter condition as point of reference when comparing *G. sulfurreducens* and *E. coli*.

**Figure 1.**
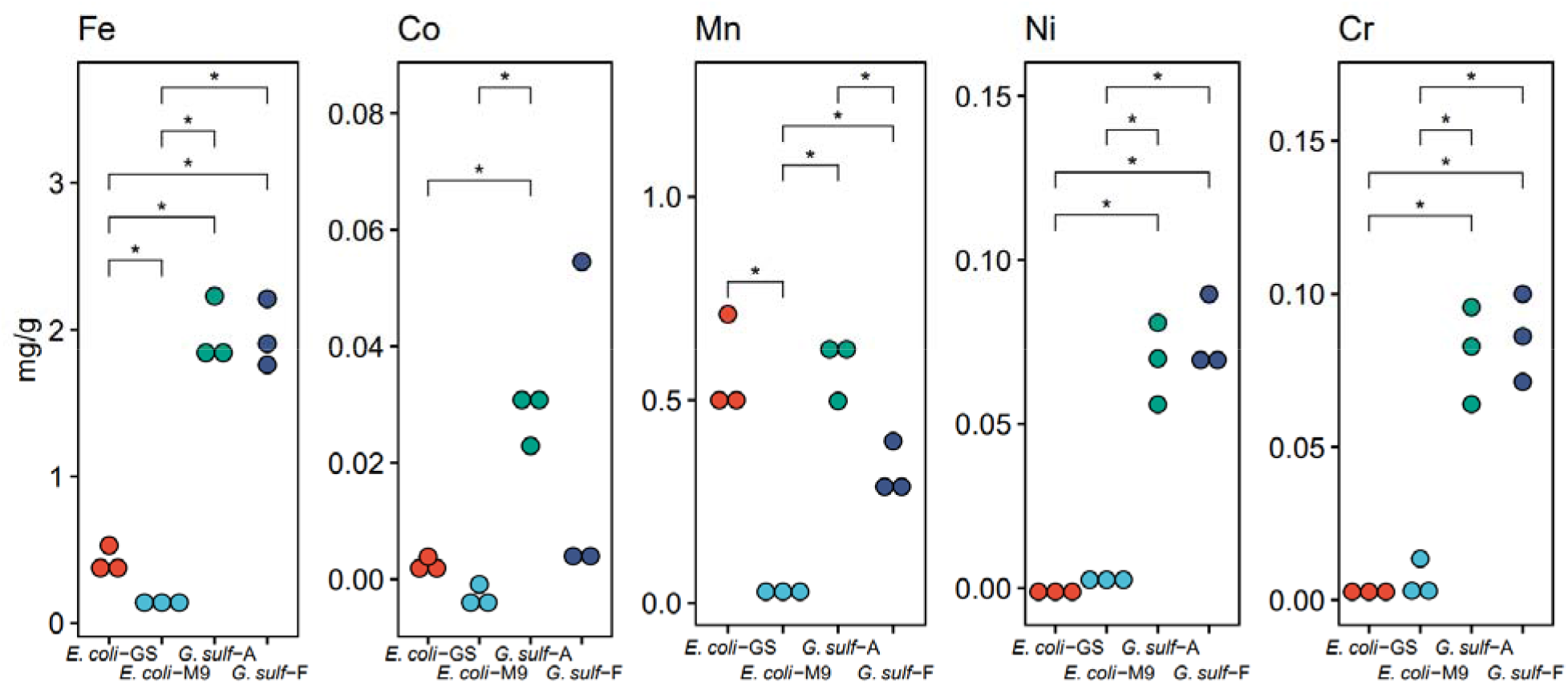
Relevant differences in metal concentrations between *E. coli-GS* (Geobacter medium), *E. coli-M9* (M9 medium), *G. sulf-A* (biofilm grown on an electrode), and *G. sulf-F* (planktonic cells using fumarate as the electron acceptor). See Figure S1 for more comparisons. *(*p*<0.05), pairwise t-test with multiple comparison correction performed with the Benjamini-Hochberg method. We chose to omit alkali metals from this figure, but Lithium did have significant differences as well (Table S1).

Several metals were present at higher concentrations in anode- and fumarate-grown *G. sulfurreducens* when compared to *E. coli*, but only a subset of them had statistically significant differences between the organisms (Table S3, Figure 1).

### Metals used as cofactors

Fe is a required trace metal for cellular respiration in many organisms as the cofactor in cytochromes. In both growth conditions for *G. sulfurreducens*, Fe was much higher than in *E. coli*, which suggest a much higher abundance of Fe-containing metalloproteins and other iron-containing biomolecules (Figure 1).

In previous studies, listed in Table 1, anaerobic bacteria are reported to have more iron per cell biomass than aerobic bacteria. Phototrophic bacteria (*Rhodospirillum*, *Rhodopseudomonas, Chromatium*) and facultative bacteria (*Escherichia, Enterobacter*) have 150-500 μg/gdw Fe. *Desulfovibrio vulgaris*, an anaerobic Deltaproteobacterium like *G. sulfurreducens*, has over 900 μg/gdw. Not included in our analysis are iron oxidizing bacteria, whose Fe precipitates can lead to Fe concentrations over 2% by dry mass (41). To our knowledge, *G. sulfurreducens* has the highest Fe content among bacteria studied, with almost twice the content of *D. vulgaris*. This is consistent with their production of heme-containing cytochromes in much higher abundance than other microorganisms, leading to not only cytoplasmic and membrane metalloproteins, but also an extensive abundance of extracellular cytochromes. Growing *G. sulfurreducens* on the anode versus fumarate did not change the total Fe amount, suggesting similar abundance of Fe metalloproteins. OmcS nanowires have been isolated from fumarate culture (6), indicating that *G. sulfurreducens* does not necessarily downregulate its EET metabolism when it is not needed. Our gene expression data also shows a high expression of cytochromes associated with EET in fumarate and anode biofilm culture (Figure 2).

**Table 1.**
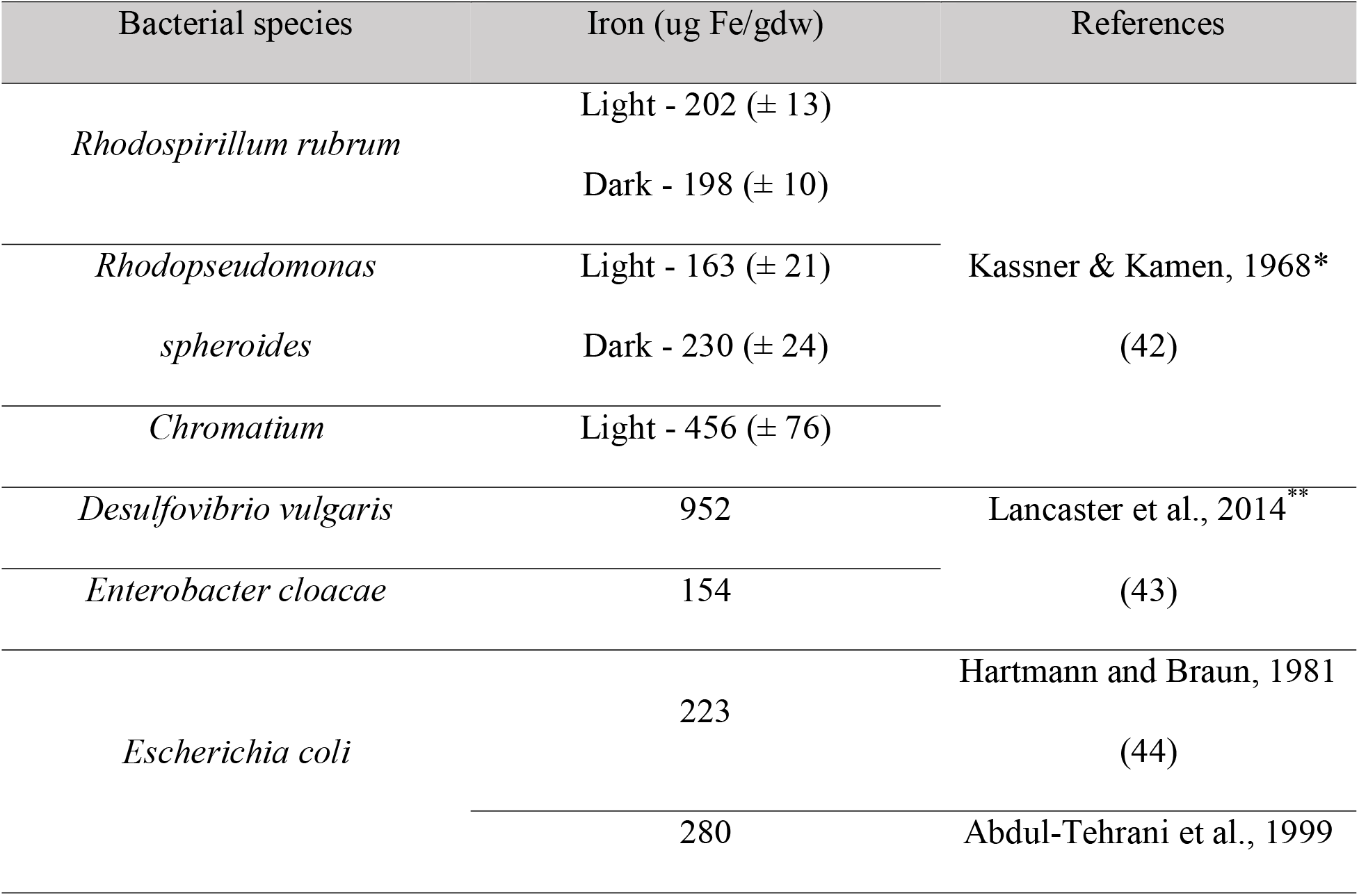
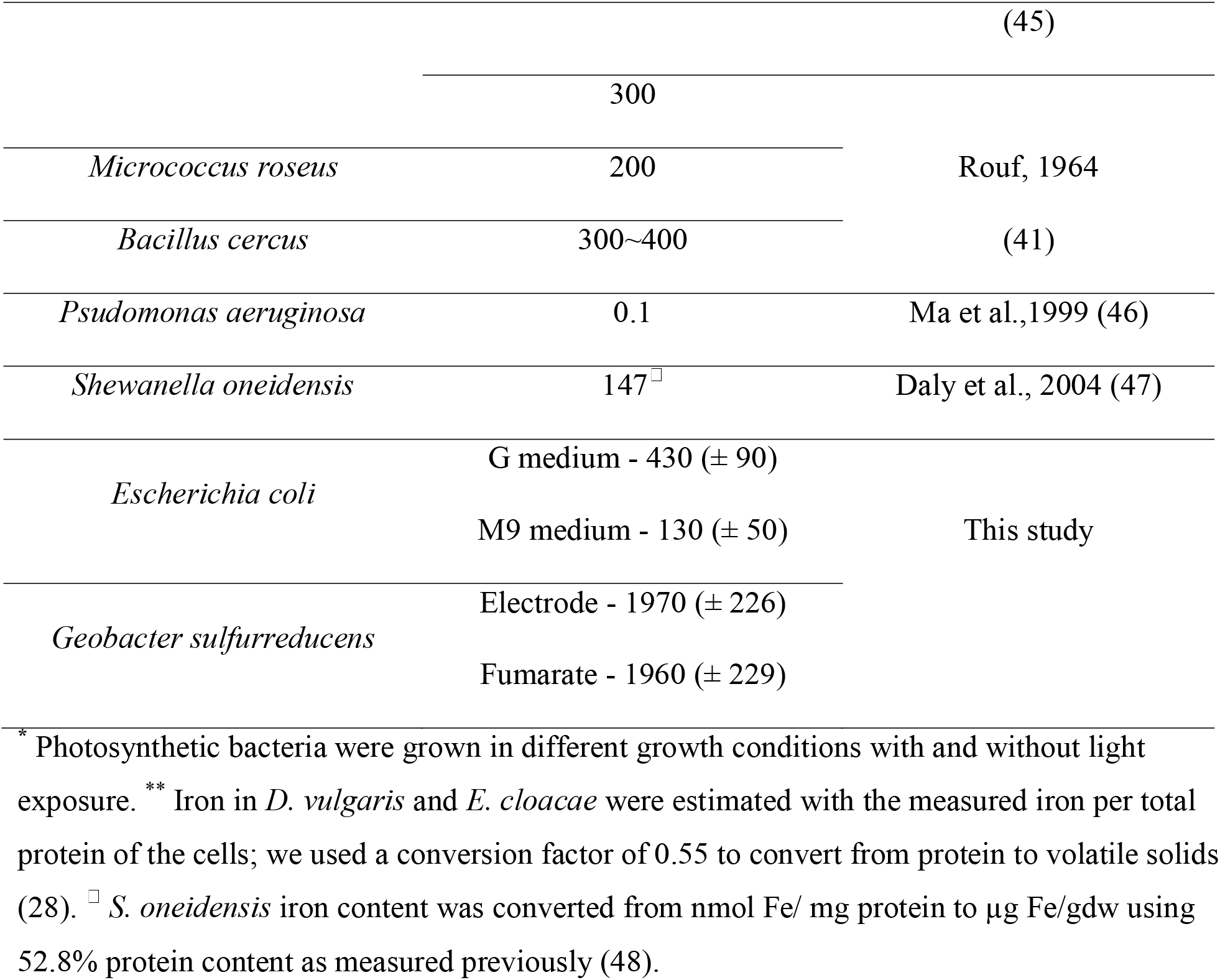
Amount of iron in different prokaryotic species

**Figure 2.**
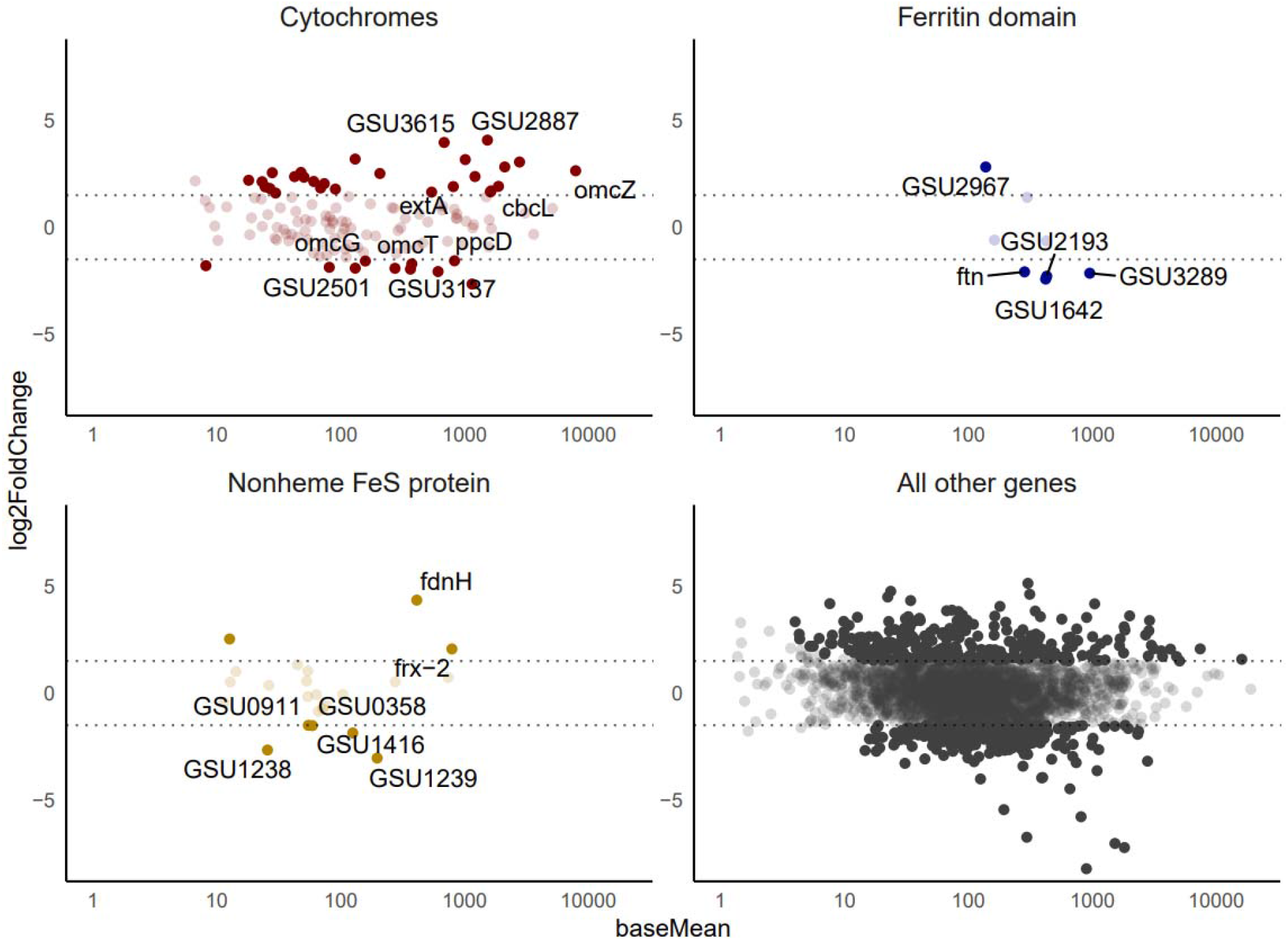
MA plots of mRNA gene expression data comparing planktonic cells and anode biofilms. Positive log2 fold change indicates higher expression in the anode biofilm condition. Solid points indicate a log2 fold change greater than 1.5 and an adjusted *p*-value below 0.05.

Fe may be a limiting trace mineral in commonly used *G. sulfurreducens* media. Estevez-Canales et al. found that a medium concentration of 2 μM Fe limits biomass culture in a chemostat growing *G. sulfurreducens* led to a culture with 1.9 × 10^-6^ ng of Fe per cell (15). If we assume the average cell dry weight of *G. sulfurreducens* is between 0.1 and 1 picograms, as has been shown in *E. coli* (49), our results would give an iron content of 2 × 10^-7^ ng to 2 × 10^-6^ ng Fe per cell in *G. sulfurreducens*.

Nickel, cobalt, and chrome content were significantly higher in both *G. sulfurreducens* conditions relative to *E. coli* (Table S3, Figure 1). Nickel is a cofactor in Ni-Fe hydrogenases, and the genome of *G. sulfurreducens* encodes for several (50, 51). *G. sulfurreducens* is able to assimilate cobalt through its cobamide-synthesis pathways (52), but it may also be precipitated on the cell surface as a defense mechanism against cobalt toxicity (53).

### Precipitating metals

There are some metals that may be overrepresented in our *G. sulfurreducens* samples due to precipitation. *G. sulfurreducens* requires at least two multicopper proteins, OmpC and OmpB, to respire Fe (III) oxide (54), and while these and other metalloproteins are a likely reservoir of Cu in our samples, *G. sulfurreducens* is capable of reducing Cu(II) to Cu_2_S nanoparticles that associate with cells (55). This phenomenon makes it difficult to estimate how much copper was required for metalloproteins, and how much may have been trapped as inorganic precipitates. *G. sulfurreducens* can also immobilize copper through dissimilatory reduction (56). In our data, the *G. sulfurreducens* samples were enriched in Cr compared to *E. coli* (Figure 1), while differences in Cu were not statistically significant. Manganese was significantly lower in the *E. coli* grown with M9 medium compared to all other conditions including *E. coli* grown with the *G. sulfurreducens* medium recipe because M9 medium does not contain manganese (Figure 1, Table S2).

Based on the metal content of the *G. sulfurreducens* cells collected, we can estimate a maximum cell density from the available mineral content in the common Geobacter medium (ATCC 1957). Table S4 shows the estimated growth cell assuming cells require the observed metal concentrations and only have the medium as a source. As expected, Fe is the most limiting metal in the medium, allowing for only 0.10 g cells/L. Cu and Zn are also close to this limitation and could lead to a multi-nutrient limitation when growing *G. sulfurreducens* at ~0.1 g/L. This nutrient limitation can either limit cell density in cell suspensions or limit current generation in microbial electrochemical technologies when operated in batch mode. Assuming a current production of ~0.28 A/g protein (57) or 0.6 A/g cell (based on Table 2), one liter of *Geobacter* medium can support enough *G. sulfurreducens* cells to produce 60 mA, an amount of current that is enough for most experimental setups but might be limiting in electrochemical cells with a high specific surface area.

**Table 2.**
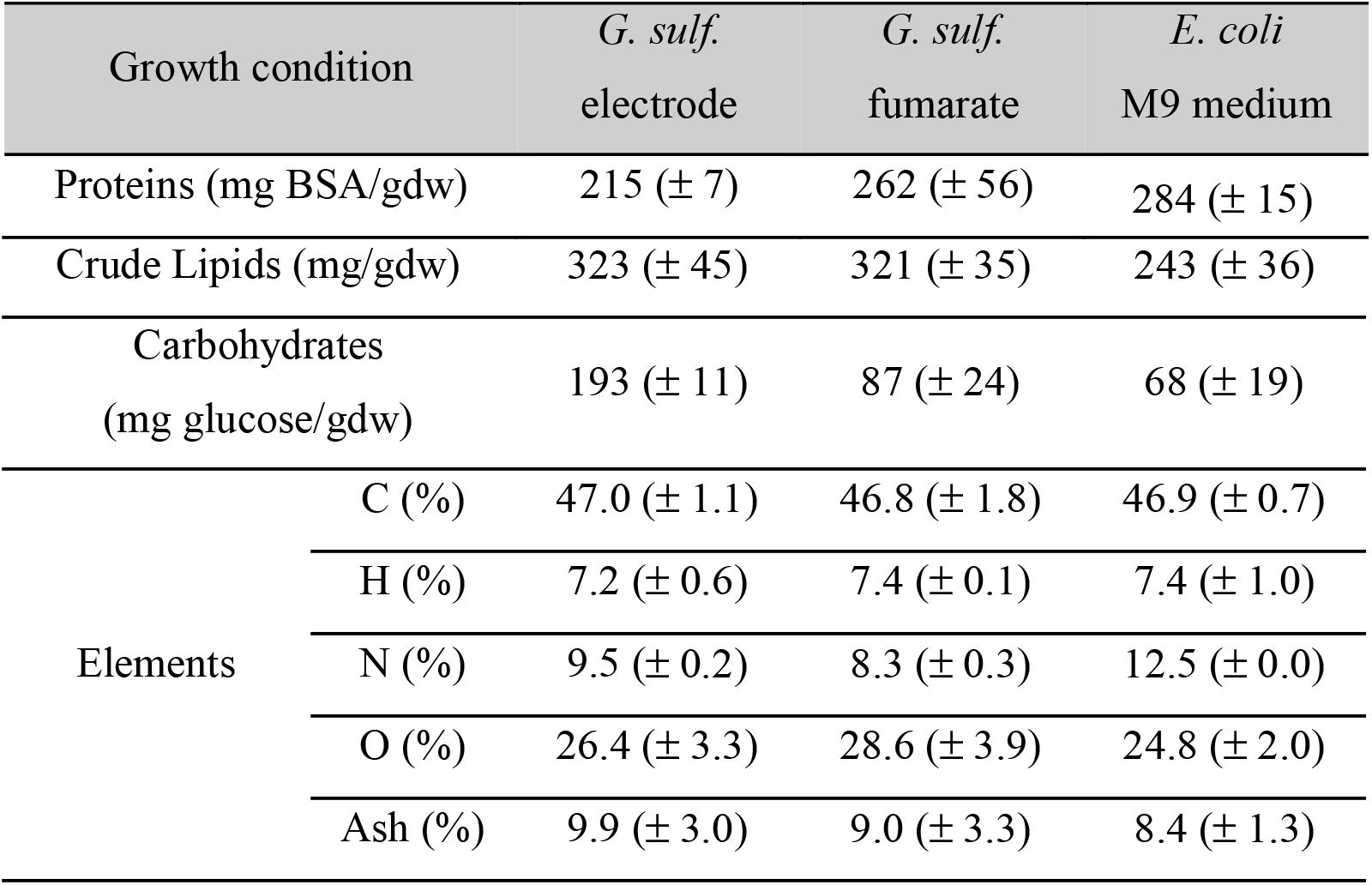
Cell compositions of *G. sulfurreducens* and *E. coli* in different growth conditions. Error is the sample standard deviation.

| Growth condition | *G. sulf.*<br>electrode | *G. sulf.*<br>fumarate | *E. coli*<br>M9 medium |
| --- | --- | --- | --- |
| Proteins (mg BSA/gdw) | 215 (± 7) | 262 (± 56) | 284 (± 15) |

|  |  |  |  |  |
| --- | --- | --- | --- | --- |
| Crude Lipids (mg/gdw) | | 323 (± 45) | 321 (± 35) | 243 (± 36) |
| Carbohydrates<br>(mg glucose/gdw) | | 193 (± 11) | 87 (± 24) | 68 (± 19) |
| Elements | C (%) | 47.0 (± 1.1) | 46.8 (± 1.8) | 46.9 (± 0.7) |
| | H (%) | 7.2 (± 0.6) | 7.4 (± 0.1) | 7.4 (± 1.0) |
| | N (%) | 9.5 (± 0.2) | 8.3 (± 0.3) | 12.5 (± 0.0) |
| | O (%) | 26.4 (± 3.3) | 28.6 (± 3.9) | 24.8 (± 2.0) |
| | Ash (%) | 9.9 (± 3.0) | 9.0 (± 3.3) | 8.4 (± 1.3) |

### Cell composition of G. sulfurreducens is different to an average bacterium

We also studied the cell composition and elemental analysis (C, H, O, N, Ash) of *G. sulfurreducens*. Interestingly, the *G. sulfurreducens* cell showed a high abundance of lipids in both growth conditions (Table 3). The values of ~32% lipid content were much higher than previously reported and similar to lipid-accumulating algal cultures (58–60). We do not know the reason why *G. sulfurreducens* requires such a high lipid content. Their smaller diameter (~ 0.5 μm) and distinct morphology (61) compared to other rod-shape bacteria certainly plays a role in the increased lipid content. *Shewanella oneidensis*, another electrogenic organism, is known to produce outer membrane extensions for electron transfer (13). While it is likely that our analysis captured some extracellular polymeric substances, the extracellular matrix of *G. sulfurreducens* has not been found to have a significant lipid component (62). Most microorganisms exhibiting this high lipid fraction have either lipid accumulation, as in the case of certain algal species (58–60) or have internal lipid structures that increase its relative fraction as in the case of thylakoid membranes and intracytoplasmic membranes (63–65). Our method for lipid quantification would likely collect other nonpolar compounds like polyhydroxyalkanoates if they were present in the biofilm, and while *G. sulfurreducens* has not been shown to produce this type of carbon storage polymer, biofilms enriched in other *Geobacter* species do use some uncharacterized carbon storage mechanism (66). We anticipate that future studies may add context to the data we present. The lipid fraction in *E. coli* was lower than *G. sulfurreducens* and higher than has been previously reported at 24.3 ± 3.6%. However, there is a high variance in reported *E. coli* lipid content with values ranging from 9 to 19% by dry weight (67, 68).

Because of the higher lipid content, *G. sulfurreducens* cells show a significantly lower protein content when compared to other microorganisms (~22-26%, Table 3). Fumarate-grown cells had a larger protein fraction than in anode-grown cells. On the other hand, total carbohydrates were ~2 times higher in anode-grown cell; exopolysaccharide (EPS) excreted from *G. sulfurreducens* to form a biofilm on the electrode probably increases the carbohydrate content in this growth condition.

Our elemental analysis of *G. sulfurreducens* cells is consistent with the low protein, high lipid content measured. Following Eq. 1, empirical cell biomass formulas of *G. sulfurreducens*,normalized to N, were calculated as C_5.77_H_10.61_O_2.43_N for electrode-grown and C_6.58_H_12.48_O_3.02_N for fumarate-grown cells. Compared with general formulas for bacterial biomass, such as C_5_H_7_O_2_N (39), *G. sulfurreducens* has a higher C:N ratio typical of a low protein content. It also has a higher hydrogen content, due to the higher lipid content that has approximately a 1:2 for fumarate-grown cells.

We compared our cellular composition of *G. sulfurreducens* to that reported in Mahadevan et al. 2006 (27). The main differences between the fraction distributions reported here and those reported in Mahadevan et al. is the higher lipid content at the expense of a lower protein content. We do not know the reason for the discrepancy, but in both cases the lipid content is significantly higher than *E. coli* and other bacterial cells.

### Iron-containing genes are highly expressed

Our analysis identified 434 genes that were differentially expressed between the anode biofilm samples and the planktonic fumarate cells out of 3434 annotated genes detected at quantifiable levels. 205 genes were more highly expressed in the anode biofilm, and 229 genes were more highly expressed in the planktonic samples. In Figure 2, MA plots visualize the differential expression and highlight several types of iron-containing protein-coding genes. While a greater number of cytochromes were significantly upregulated in the anode biofilm than the number upregulated in the planktonic samples, most cytochromes were not differentially expressed. Ferritin domain containing protein- and nonheme Fe-S domain protein-coding genes were also present among the differentially expressed genes. Our data show that iron-containing protein-coding genes are expressed in both planktonic fumarate cultures and anode biofilms, but that there are specific iron-related genes whose expression depends on growth conditions. The abundance of expression of Fe-containing proteins is consistent with the high Fe abundance in both conditions. Our gene expression data is similar in overall patterns to the data in Otero et al. 2018, with multiheme cytochromes like the PpcA, OmcZ, and CbcL among the most expressed genes overall, and only a subset of all cytochromes differentially expressed between electrode and fumarate conditions (7). We did, however, observe a greater number of cytochrome genes differentially expressed between electrode and fumarate conditions than Otero et al. 2018, likely due to the lower electrode potential that we used. We also investigated the expression of lipid synthesis pathway genes and found that many of them were present in the transcriptome but not differentially expressed, although a few lipid-associated genes showed significant differences (Figure S2).

## Conclusions

*G. sulfurreducens* is a bacterium with a complex system of electroactive proteins, and those electroactive proteins largely require iron. This may be a factor in the high iron concentration we measured in *G. sulfurreducens* relative to non-electroactive Gram-negative *E. coli* and values reported in literature for other bacteria. Our analysis complements previous work showing that restricting iron limits EET in *G. sulfurreducens* (15). This study estimates what the nutrient limitations might be for *G. sulfurreducens*, and this information is valuable for biotechnologists developing applications using this and similar organisms. The nearly identical composition between anode-grown and fumarate-grown cells supports the hypothesis that *G. sulfurreducens* is not adapted to efficiently grow on fumarate – it makes electron carriers for EET regardless of the electron acceptor if the nutrients are available. The lipid content measured in *G. sulfurreducens* was higher than what has been reported before, and relatively high for a bacterium without lipidic storage. While all samples were taken from active biofilms or suspended cultures, we did not have a mechanism to separate dead cells from active cells, and it is probable that the composition of an individual cell may differ from the composition of the bulk samples analyzed. When compared to similar studies on other bacteria and the *E. coli* in our study, we have shown that *G. sulfurreducens* has a unique composition to support its complex metabolism.

## Acknowledgements

The funding for this work was provided by Office of Naval Research (ONR awards N0014-15-1-2702 and N0014-20-1-2269). We also thank Roy Erickson and Adam Smith for assistance of ICP-OES and elemental analysis (Goldwater Environmental Laboratory at ASU) and the Genomics Core at ASU for sequencing.

## Notes

### Competing Interest Statement

The authors have declared no competing interest.

### Summary of Updates

Figure 2 revised; supplemental file updated, added additional discussion about lipid genes; updated lipid section to mention polyhydroxyalkanoates, updated estimate of the mass of a single cell; added a reference to compare our data to other E. coli data; amended gene expression section to discuss additional cytochromes and references

