## Supplemental Information 1 for "Geobacter sulfurreducens’ unique metabolism results in cells with a high iron and lipid content"

**Supporting Information**

**Table S1**. Chemicals in Ringer's solution per L of medium used. One fourth strength of Ringer's solution was used for washing cells.

| Chemicals | Amount (g) | Concentration (mM) | ¼ Strength Ringer's solution Conc. (mM) |
|---|---|---|---|
| NaCl | 6.5 | 111 | 27.8 |
| KCl | 0.42 | 5.6 | 1.41 |
| CaCl2 | 0.25 | 2.3 | 0.56 |
| NaHCO3 | 0.2 | 2.4 | 0.6 |

**Table S2. Media composition in different conditions tested.**

| Species | *Geobacter sulfurreducens* PCA | | | | *Escherichia coli* K-12 | | | |
|---|---|---|---|---|---|---|---|---|
| Growth condition | Electrode | | Fumarate | | G medium | | M9 medium | |
| Unit | g in 1 L | mM | g in 1 L | mM | g in 1 L | mM | g in 1 L | mM |
| $NaCH_3COO$ $3H_2O$ | 6.8 | 50 | 6.8 | 50 | - | - | - | - |
| NaFumarate | - | - | 16 | 100 | - | - | - | - |
| Glucose | - | - | - | - | 8 | 44 | 8 | 44 |
| $NaHCO_3$ | 2.5 | 30 | 2.5 | 30 | 2.5 | 30 | - | - |
| $NaH_2PO_4$ | 0.6 | 4 | 0.6 | 4 | 0.6 | 4 | 25.6 | 180 |
| $KH_2PO_4$ | - | - | - | - | - | - | 6 | 44 |
| KCl | 0.1 | 1 | 0.1 | 1 | 0.1 | 1 | - | - |
| NaCl | - | - | - | - | - | - | 1 | 17 |
| $NH_4Cl$ | 1.5 | 20 | 1.5 | 20 | 1.5 | 20 | 2 | 37 |
| $MgSO_4$ $7H_2O$ | - | - | - | - | - | - | 0.049 | 0.200 |
| $CaCl_2$ | - | - | - | - | - | - | 0.001 | 0.010 |
| Thiamin | - | - | - | - | - | - | 0.001 | 0.024 |
| $FeCl_2$ $4H_2O$ | 0.004 | 0.020 | 0.004 | 0.020 | 0.004 | 0.020 | - | - |
| $Na_2S$ $9H_2O$ | 0.013 | 0.054 | 0.013 | 0.054 | 0.013 | 0.054 | - | - |
| vitamin mix | 10 mL | 10 mL | 10 mL | 10 mL | 10 mL | 10 mL | - | - |
| trace mineral* | 10 mL | 10 mL | 10 mL | 10 mL | 10 mL | 10 mL | - | - |

* Trace mineral stock solution in 1 L

| Trace mineral | Al | B | Ca | Cl | Co | Cu | Fe* | K* |
|---|---|---|---|---|---|---|---|---|
| mg in 1 L | 0.006 | 0.014 | 0.273 | 7.595 | 0.248 | 0.025 | 0.201 | 0.008 |
| Trace mineral | Mg | Mn | Mo | Na* | Ni | S* | W | Zn |
| mg in 1 L | 2.958 | 1.625 | 0.107 | 3.989 | 0.059 | 5.656 | 0.014 | 0.624 |

**Table S3**. Metal content in dry cell mass of *Geobacter sulfurreducens* PCA and *Escherichia coli* K-12 (unit: milligram of each metal per gram of dried bacterial cell, mg/gdw).

| | ***G. sulfurreducens*** | | ***E. coli*** | |
|---|---|---|---|---|
| | Anode | Fumarate | M9 medium | *Geobacter* medium |
| Ag | 0.098 (± 0.073) | 0.106 (± 0.034) | ND | ND |
| Ba | 0.028 (± 0.002) | 0.025 (± 0.003) | 0.021 (± 0.005) | 0.027 (± 0.003) |
| Co | 0.028 (± 0.005) | 0.021 (± 0.029) | ND | 0.003 (± 0.002) |
| Cr | 0.081 (± 0.016) | 0.086 (± 0.014) | 0.006 (± 0.003) | 0.003 (± 0.000) |
| Cu | 0.224 (± 0.086) | 0.471 (± 0.107) | 0.017 (± 0.005) | 0.048 (± 0.010) |
| Fe | 1.970 (± 0.226) | 1.96 (± 0.229) | 0.134 (± 0.053) | 0.428(± 0.089) |
| Li | 0.047 (± 0.030) | 0.017 (± 0.001) | 0.040 (± 0.003) | 0.047 (± 0.006) |
| Mg | 0.67 (± 0.27) | 0.54 (± 0.11) | 2.07 (± 0.75) | 1.79 (± 0.38) |
| Mn | 0.583 (± 0.076) | 0.324 (± 0.067) | 0.023 (± 0.021) | 0.570 (± 0.122) |
| Ni | 0.069 (± 0.012) | 0.076 (± 0.012) | 0.002 (± 0.003) | ND |
| Pb | 0.036 (± 0.015) | 0.023 (± 0.003) | 0.018 (± 0.011) | 0.016 (± 0.010) |
| Se | 0.104 (± 0.058) | 0.177 (± 0.112) | 0.012 (± 0.005) | 0.101 (± 0.016) |
| Sr | 0.008 (± 0.002) | 0.017 (± 0.006) | 0.005 (± 0.003) | 0.003 (± 0.001) |
| V | 0.002 (± 0.001) | 0.003 (± 0.002) | 0.035 (± 0.006) | 0.045 (± 0.005) |
| Zn | 3.5 (± 1.9) | 10.0 (± 2.4) | 0.135 (± 0.034) | 0.176 (± 0.024) |

**Table S4.** Estimated growth of *G. sulfurreducens* per L of medium at an anode based on the available mineral concentrations in Geobacter medium.

| Metal | *Geobacter* Medium [mg element/L] | Average *G. sulfurreducens* content (mg element/g cell) | Estimated growth (g cells/L) |
|---|---|---|---|

| | | | |
|---|---|---|---|
| Mg | 2.958 | 0.67 | 4.42 |
| Mn | 1.625 | 0.583 | 2.79 |
| Fe | 0.201 | 1.97 | 0.10 |
| Co | 0.248 | 0.028 | 8.80 |
| Zn | 0.624 | 3.5 | 0.18 |
| Cu | 0.025 | 0.224 | 0.11 |
| Ni | 0.059 | 0.069 | 0.86 |

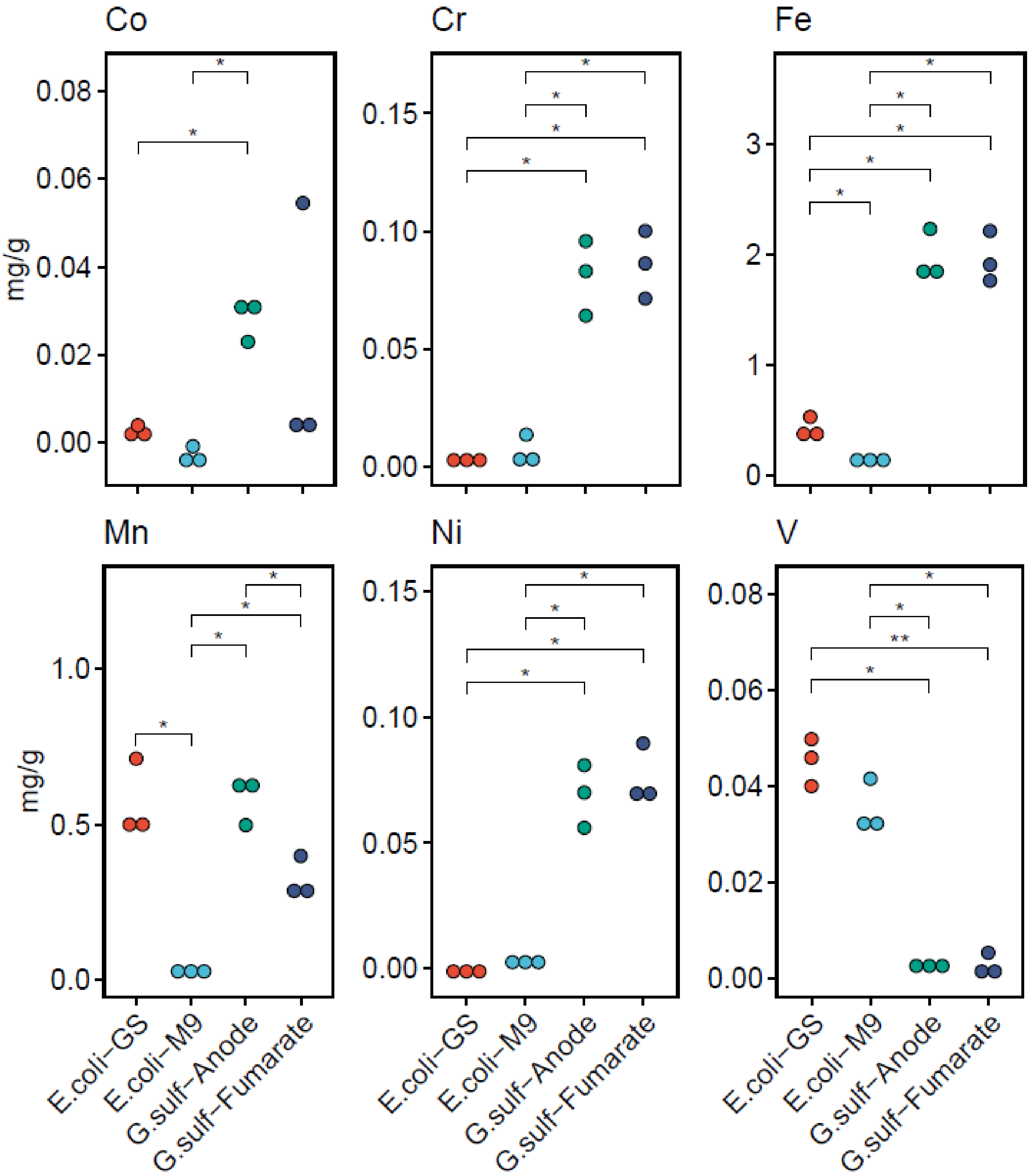


**Figure S1**. Statistically significant differences in metal concentrations between *E. coli*-GS (Geobacter medium), *E. coli*-M9 (M9 medium), *G. sulf*-Anode (biofilm grown on an electrode), and *G. sulf*-Fumarate (planktonic cells using fumarate as the electron acceptor). *($p$<0.05), **($p$<0.001), pairwise t-test with multiple comparison correction performed with the Benjamini-

Hochberg method. We chose to omit alkali metals from this figure, but Lithium did have significant differences as well (Table S3).

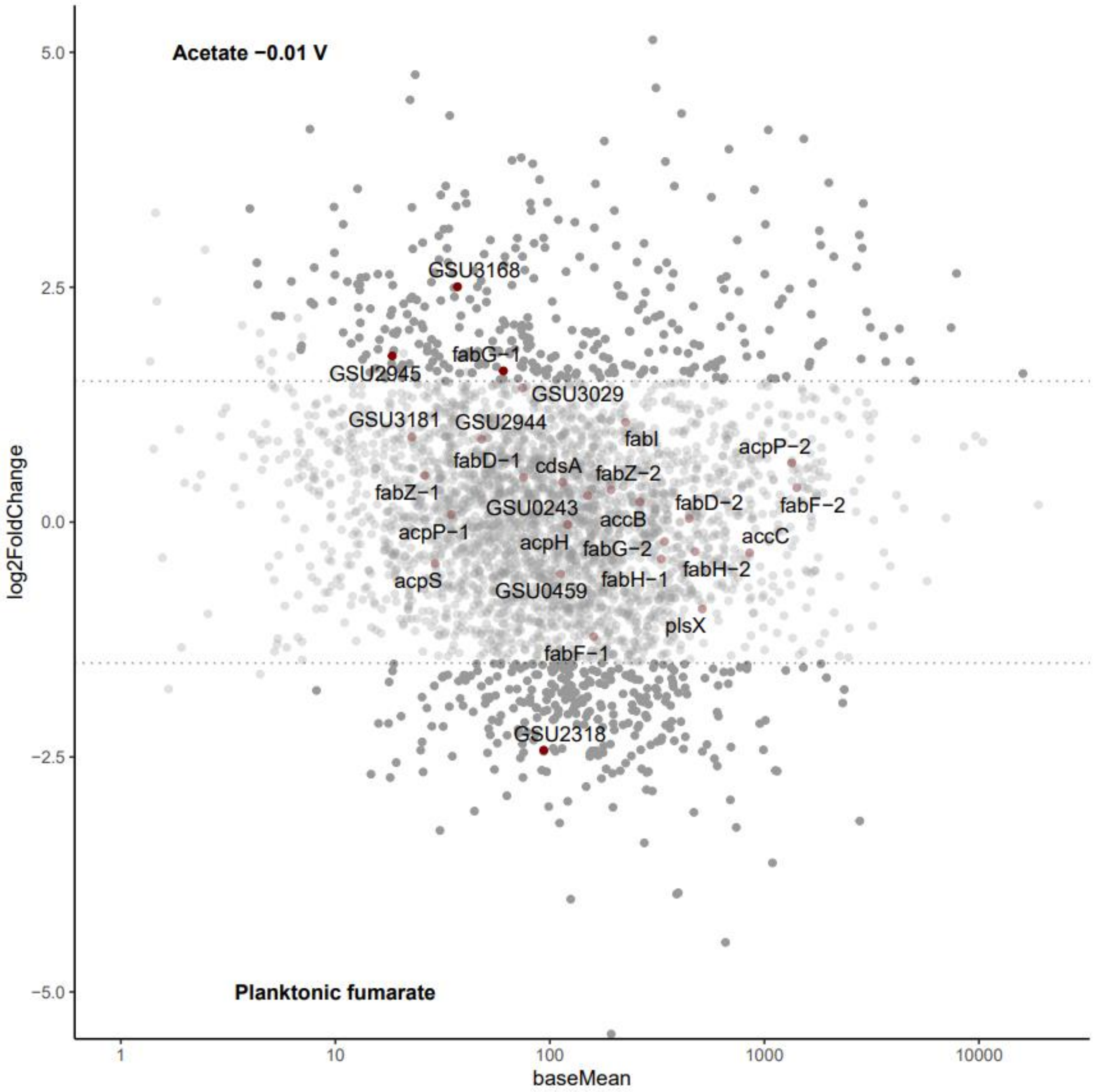


**Figure S2**: MA plot with lipid synthesis pathway genes annotated. Dotted lines indicate $\log_2$ fold change of 1.5, and solid dots indicate an adjusted p value under 0.05.
